# Run-on sequencing reveals nascent transcriptomics of the human microbiome

**DOI:** 10.1101/2022.04.22.489220

**Authors:** Albert C. Vill, Edward J. Rice, Iwijn De Vlaminck, Charles G. Danko, Ilana L. Brito

## Abstract

Precise regulation of transcription initiation and elongation enables bacteria to control cellular responses to environmental stimuli. RNAseq is the most common tool for measuring the transcriptional output of bacteria, comprising predominantly mature transcripts. To gain further insight into transcriptional dynamics, it is necessary to discriminate actively transcribed loci from those represented in the total RNA pool. One solution is to capture RNA polymerase (RNAP) in the act of transcription, but current methods are restricted to culturable and genetically tractable organisms. Here, we apply precision run-on sequencing (PRO-seq) to profile nascent transcription, a method amenable to diverse species. We find that PRO-seq is well-suited to profile small, structured, or post-transcriptionally modified RNAs, which are often excluded from RNAseq libraries. When PRO-seq is applied to the human microbiome, we identify taxon-specific RNAP pause motifs. We also uncover concurrent transcription and cleavage of guide RNAs and tRNA fragments at active CRISPR and tRNA loci. We demonstrate the specific utility of PRO-seq as a tool for exploring transcriptional dynamics in diverse microbial communities.

## INTRODUCTION

Bacterial transcriptional circuitry underlies cellular stress responses, host-pathogen immune interactions, group-level dynamics, and other responses to environmental stimuli. Within the gut microbiome, these transcriptional responses may reveal pathways involved in pathogenesis or define the resilience of communities under different selective pressures. Metagenomic sequencing has been used to infer the potential functions of microbiome constituents, though only a fraction of genes in a cell are expressed at any given time. RNAseq has therefore been used to provide a more accurate depiction of cellular function. However, RNAseq, as performed on microbiomes, gives limited information about transcriptional dynamics across genes, requires depletion of ribosomal RNA, which may introduce species- and sequence-specific biases, and may fail to capture small, structured, or post-transcriptionally modified RNAs.

RNAseq indiscriminately sequences the pool of mature and accessible RNA molecules. In comparison, the nascent transcriptome comprises only RNA molecules that are being actively transcribed by RNA polymerase (RNAP). While total RNA sequencing has great utility in measuring steady-state levels of messenger RNA, the nascent transcriptome represents the state of a cell agnostic to the different degradation rates of RNA species. In model eukaryotes, nascent transcriptomics has aided the study of RNAP kinetics and revealed species of transient noncoding RNAs important for transcriptional regulation (reviewed in ^1^).

In bacteria, nascent transcriptomics has shed light on the pausing and elongation dynamics of RNAP. However, these observations have been largely limited to genetically tractable model organisms due to significant methodological constraints. NET-seq involves the immunoprecipitation of RNAP, and thus requires either clade-specific RNAP antibodies or genetic manipulation to add epitope tags; to date, it has only been applied to *Escherichia coli* and *Bacillus subtilis* ^2,3^. Other methods rely on discrimination of mature and immature RNAs by enzymatic recognition of 5’ nucleotide chemistry. Differential RNA-seq (dRNA-seq) has been applied to diverse bacterial species and employs 5’-P-dependent exonuclease to degrade monophosphorylated mature transcripts, leaving immature triphosphorylated transcripts to be sequenced ^4–6^. Likewise, Cappable-seq has been applied to *E. coli* and a mouse cecal microbiome and relies on a 5’-PPP capping enzyme to incorporate biotin into nascent transcripts in order to map transcription start-sites (TSS) ^7^. While these methods are well-equipped to identify TSSs by mapping 5’ transcript ends, they do not provide information about the position and procession of RNAP.

Precision run-on sequencing (PRO-seq) has been developed to uncover transient transcriptional signals in eukaryotes ^8–10^. PRO-seq involves capturing RNA bound by engaged and actively transcribing RNAP (Figure 1A). In PRO-seq, cells are first permeabilized to deplete endogenous nucleotide triphosphates (NTPs), halting transcription. Then, lysates are subject to a ‘run-on’ reaction, which introduces biotinylated NTPs to reinitiate transcription and tag the 3’ ends of nascent transcripts. Nascent RNA molecules are then enriched using streptavidin-coated beads and sequenced. Apart from eukaryotic RNAPII, transcription elongation by run-on reaction has been demonstrated for T7 RNAP ^11^ and mitochondrial POLRMT ^12,13^, suggesting that PRO-seq may be amenable to RNA polymerases across the tree of life. Here, we establish PRO-seq as a method for prokaryotes using an *E. coli* heat shock model and in human gut microbiomes, to measure nascent transcription across diverse species simultaneously.

**Figure 1.**
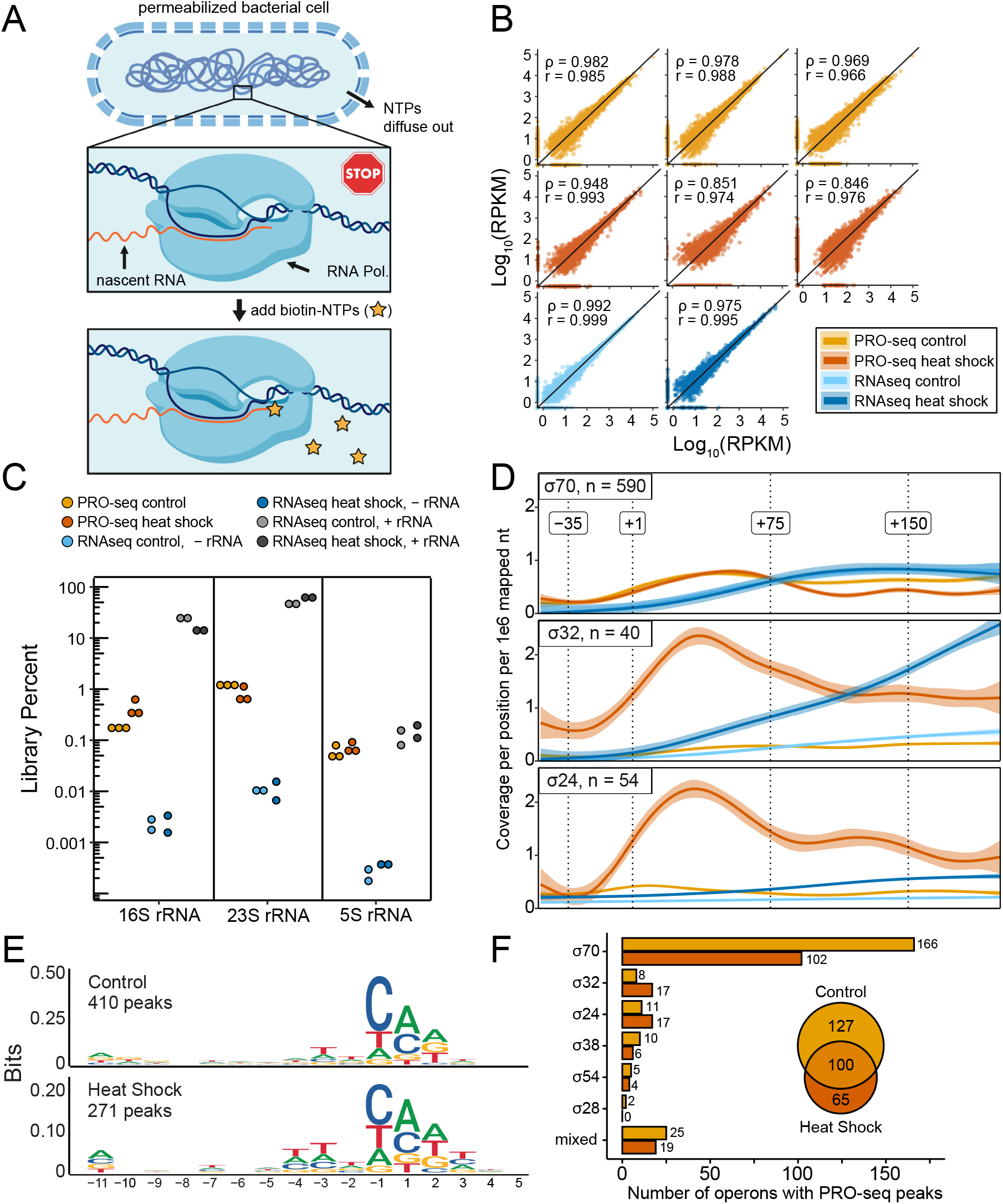
PRO-seq captures nascent transcripts in *E. coli*. (A) Outline of bacterial PRO-seq. Cells are permeabilized to liberate NTPs and halt RNA polymerization. Addition of biotinylated NTPs allows RNA polymerase to incorporate a single biotin-NTP to the 3’ end of the nascent RNA strand. (made with BioRender) (B) Correlation of reads aligning to genes in *E. coli* in replicate samples in rRNA-depleted RNAseq (n = 2) and PRO-seq (n = 3) libraries in control and heat shock-treated *E. coli* cells. Spearman’s rank correlation coefficients (*ρ*) and Pearson’s correlation coefficients (*r*) are inset. (C) Percent of reads aligning to *E. coli* 16S, 23S and 5S rRNA genes in RNAseq libraries without rRNA depletion, RNAseq libraries with rRNA depletion, and PRO-seq libraries made from control and heat shock-treated samples. (D) Normalized and smoothed mean read depth profiles proximal to *E. coli* transcription start sites (TSS, position +1) under control of promoters regulated by σ^70^, σ^32^, and σ^24^, as annotated by RegulonDB v10.9. Replicate libraries were combined for each library type + treatment pair: rRNA-depleted RNAseq control (light blue), PRO-seq control (light orange), rRNA-depleted RNAseq heat shock (dark blue), and PRO-seq heat shock (dark orange). For RNAseq libraries, composite profiles represent full reads, whereas PRO-seq profiles only represent read 3’ ends. For operons under the control of multiple promoters, plots are centered at the TSS closest to the start codon of the first gene, and operons regulated by multiple sigma factors are excluded. Bounds represent normal confidence intervals. (E) Logos for sequences surrounding PRO-seq read 3’ end peaks coincident with regulatory regions, which are defined for each operon as the sequence starting from the left-most TSS and ending with the first base of the start codon of the first gene. The range of nucleotides in physical association with *E. coli* RNAP is plotted (−11 to +5), where position −1 represents the RNAP pause site and position 1 represents the next nucleotide added. (F) For peaks and regulatory regions described in (E), bar plots show the distribution of sigma factors regulating promoters within regulatory regions containing one or more peaks. “Mixed” regulatory regions contain promoters under control of two or more different sigma factors. The inset Venn diagram shows the overlap between peak-containing regulatory regions for control and heatshock libraries, replicates merged.

## RESULTS

### Paired PRO-seq and RNA-seq discriminate promoters by sigma factors in E. coli

The response to heat shock in *E. coli* is controlled, in part, at the level of transcription. We performed an experiment comparing RNAseq and PRO-seq in *E. coli* MG1655 cells subject to 7 minutes of heat-shock at 50 °C, hypothesizing that we could identify differences in cellular responses at genes controlled by specific sigma factors involved in the heat shock response. Pairwise scatterplots of technical triplicates show that PRO-seq is replicable in bacteria (Figure 1B), and, as expected, rank-ordering of transcripts suggests that PRO-seq and RNAseq signals are correlated (*ρ* = 0.875 for control; *ρ* = 0.76 for heat-shock, Spearman’s rank correlation). Bacterial RNAseq requires ribosomal RNA (rRNA) depletion to reduce rRNA representation in sequencing; across *E. coli* treatments, rRNA depletion reduces rRNA from 73.7 ± 2.7% to 0.013 ± 0.004% of the library. In contrast, *E. coli* PRO-seq libraries are 1.39 ± 0.31% rRNA reads, demonstrating that PRO-seq is agnostic to bias from highly stable RNA species (Figure 1C). Removing the need for rRNA depletion has the benefit of reducing the potential bias introduced by such handling steps ^14^.

Examining the RNAseq data, there was no difference between the read depth profiles proximal to transcription start sites under control of σ70 promoters in the different treatments, consistent with the role of σ70 as the major regulator of housekeeping genes (Figure 1D). This was also apparent in the PRO-seq data, where the position of RNA polymerase is denoted by the 3’ ends of nascent transcripts. During heat shock, the PRO-seq profiles across the same loci were comparatively reduced as transcription continues into gene bodies. This may be explained by aborted transcription of housekeeping genes in favor of genes needed to mount a response to thermal stress. Accordingly, at operons regulated by σ32, the master regulator of the heat shock response, we saw upregulation in both the RNAseq and PRO-seq datasets upon heat shock. The σ24 envelope stress response is only active in response to extreme heat stress^15,16^. Transcription proximal to σ24 promoters is solely captured by PRO-seq during heat shock. These data suggest that PRO-seq enables the observation of active loading of RNA polymerase at σ24-controlled loci preceding the accumulation of mature transcripts.

We were also able to identify pause site motifs in *E. coli* using PRO-seq (Figure 1E). We defined PRO-seq peaks as any genomic position centered in a 50 bp window with a minimum 3’ read end depth of 10 and a Z-score of at least 5. RNAP pause sites were found at both 5’ untranslated regions and within gene bodies, suggesting that PRO-seq can be used to uncover promoter-adjacent regulatory pausing as well as elemental pausing. The sigma factor repertoires of peak-containing regulatory regions are concordant with the treatments: heat-shocked *E. coli* operons regulated by σ32 and σ24 are enriched in the merged heat shock dataset relative to the control (Figure 1F).

### PRO-seq is suitable for diverse species of human-associated microbiota

We next investigated the utility of PRO-seq to capture nascent transcripts from diverse microbial communities. We performed PRO-seq and RNAseq, with replicates, on gut microbiome samples from two healthy individuals. The first step in performing PRO-seq is permeabilization, which results in the rapid depletion of NTPs from cells, thus halting transcription. We were concerned that the permeabilization protocol used on *E. coli* would be insufficient to permeabilize microbiome-derived cells, as harsher lysis methods are typically required to minimize extraction biases ^17,18^. As heat and proteinase treatment are incompatible with PRO-seq, we opted for bead-beating in a nonionic detergent buffer, preserving halted RNAP-RNA complexes (which can be very stable ^19^) for subsequent run-on reactions. Overall, we found strong concurrence between replicate samples (*ρ* = 0.954 and *ρ* = 0.938 for the two microbiome samples, Spearman’s rank correlation) (Figure 2A). We subset reads according to the metagenomic assembled genomes to which they aligned, and found an enrichment of certain strains in the PRO-seq libraries compared to the RNAseq libraries (Figure 2B). With the exception of *Ruminococcus bromii*, Firmicutes were less well represented in the PRO-seq libraries than RNAseq libraries. This may be attributed to more efficient lysis of Gram negative Bacteroidetes. Alternatively, Bacteroidetes are highly abundant in both of these samples, which may reflect their overall higher growth rates and possibly more active transcription.

**Figure 2.**
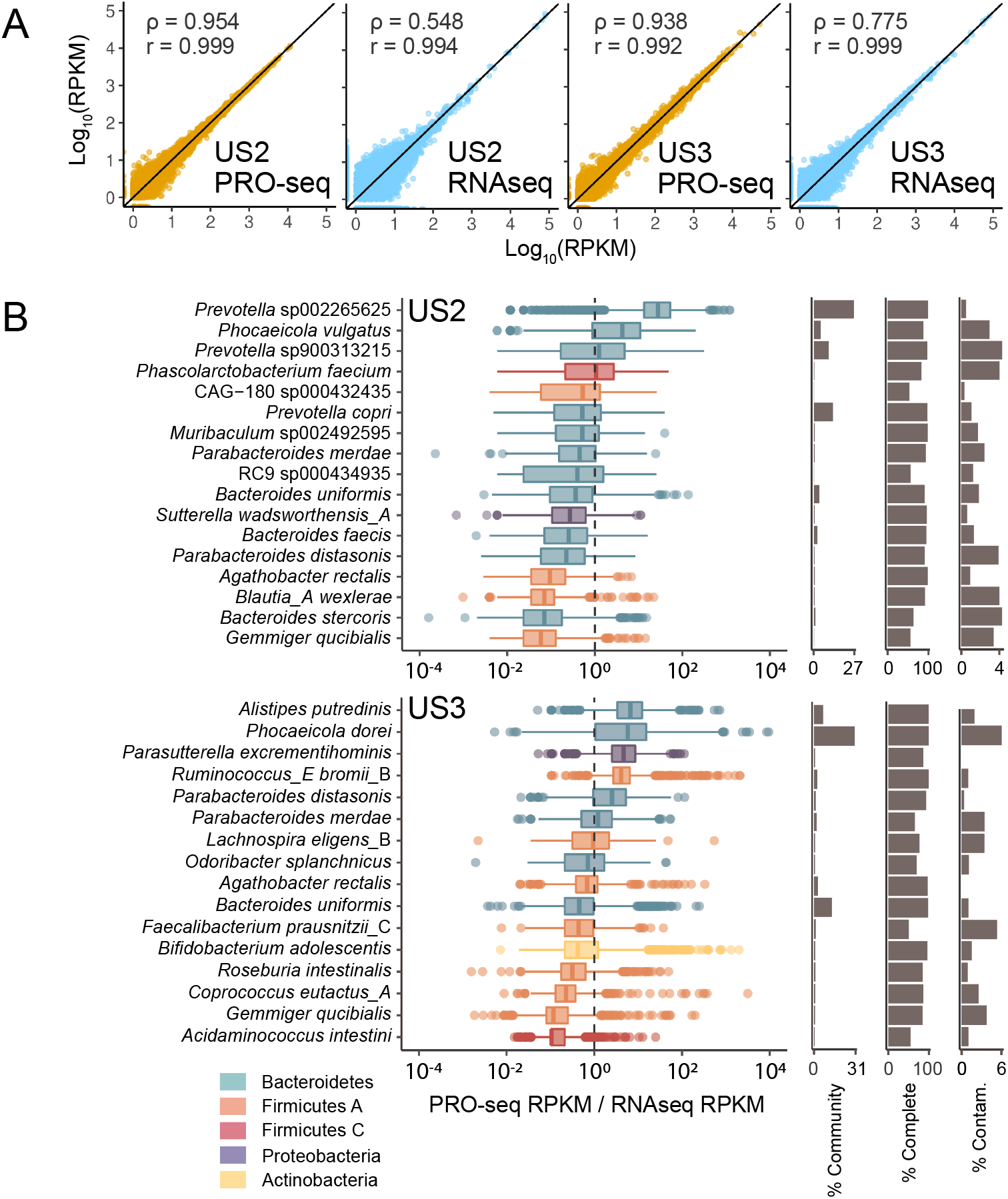
Relative coverage of bacterial species in PRO-seq samples compared to their corresponding metagenomes. (A) Correlation of reads aligning to metagenomic features for replicate samples in rRNA-depleted RNAseq (n = 2) and PRO-seq (n = 2) libraries. Spearman’s rank correlation coefficients (*ρ*) and Pearson’s correlation coefficients (*r*) are inset. (B) For metagenomic bins that are least 90% complete with less than 5% contamination, box plots show the distribution of PRO-seq RPKM divided by RNAseq RPKM for each feature; replicate libraries are merged. Dotted lines demarcate equal coverage in both sequencing types. For each bin, relative abundance, percent completeness, and percent contamination are provided (right).

### PRO-seq captures concurrent transcription and cleavage of CRISPR RNAs

We next turned our focus towards specific genomic loci that tend to be difficult to capture by RNAseq. Non-coding RNAs may be structured or sequestered in protein complexes, affecting their representation in metatranscriptomic experiments ^20^. CRISPR arrays are comprised of repeated elements and unique spacers which are transcribed and cleaved to create functional guide RNAs. RNAseq reads that align to CRISPR loci are typically sparse ^21^. Whereas CRISPR arrays are lowly represented in our RNAseq data as well, we see active transcription across these loci in the PRO-seq data. Furthermore, at CRISPR loci with high PRO-seq coverage, we observe a distinct periodic pattern with a pile-up of PRO-seq read 5’ ends at consistent positions within repeats (Figure 3A, Supplemental Figures 1A & 1B). When examining further, we found these pile-ups occur at predicted sites of endonuclease hydrolysis, corresponding to the 3’ ends of the predicted repeat stem loops (Figure 3B). It is currently unclear whether transcription of the full pre-crRNA precedes endonuclease processing or if pre-crRNAs are co-transcriptionally cleaved. Our data suggest that the latter is the case, as the capture of individual crRNAs in our PRO-seq libraries implies those transcripts are bound by RNAP. In support of this finding, on a contig for which we were able to assemble a CRISPR array and its associated Cas proteins, we find active expression of the upstream Cas5d endonuclease (Figure 3C). Given that in most CRISPR systems, the newest spacers are incorporated at the end of the array closest to the leader ^22^, this model of co-transcriptional cleavage is consistent with the need to rapidly assemble CRISPR-Cas complexes to respond to incoming phage or other mobile genetic elements.

**Figure 3.**
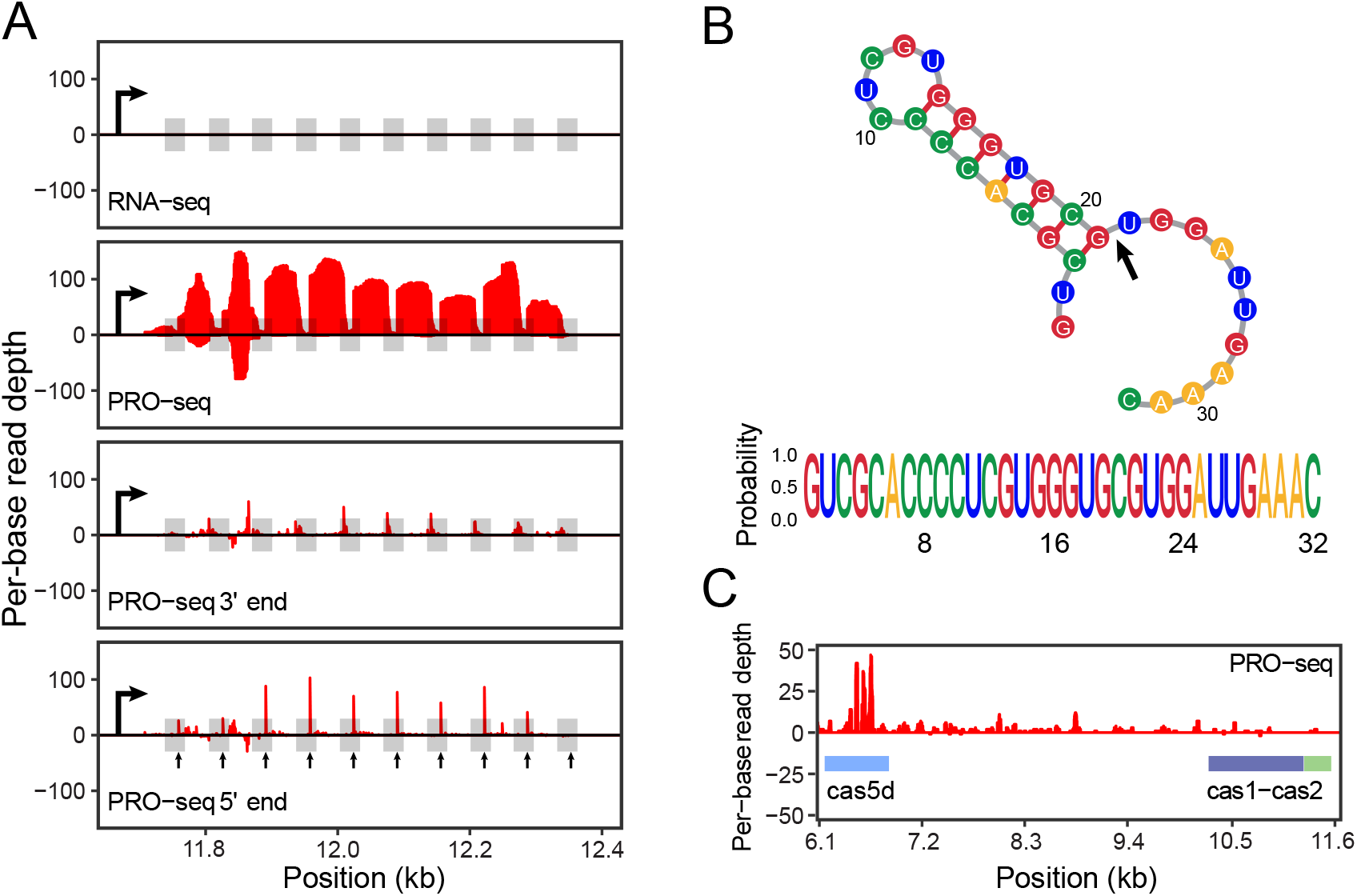
Nascent transcription of an active CRISPR loci reveals transcriptional pre-crRNA processing. (A) Coverage of PRO-seq and RNA-seq reads across a CRISPR array in a US2 *Prevotella* sp. contig. Shaded boxes represent repeats. The large black arrow in each panel represents the leader sequence containing a putative promoter. Small black arrows in the “PRO-seq 5’ end” panel correspond to the predicted site of crRNA cleavage proximal at the base of the repeat stem loop. (B) Predicted crRNA repeat secondary structure. The black arrow points to the phosphodiester bond that is likely cleaved by Cas5d during pre-crRNA processing, which marks the same position in the repeat as the small arrows in (A). The sequence logo shows perfect conservation of the repeat sequence for this array. (C) PRO-seq captures nascent transcription of cas5d upstream of and contiguous with the CRISPR array.

PRO-seq profiles indicate additional transcriptional dynamics at CRISPR loci. For instance, we detect anti-sense transcription for a subset of the spacers closest to the leader (Figure 3A), a phenomenon that has been previously observed but whose functional significance is poorly understood ^21,23–25^. In some of the detected CRISPR arrays, pile-ups of PRO-seq read 5’ ends are coincident with spacers, not repeats, indicative of pre-crRNA processing in some systems employing RNase III ^26^. This implies that PRO-seq can capture transcription across CRISPR arrays that undergo diverse modes of maturation. We also observe regular 3’ end transcript pile-ups at specific nucleotides within the CRISPR array (Figure 3A, Supplemental Figure 1), which may point to regulation at these loci at the level of RNAP procession and dissociation.

### Concurrent transcription and cleavage also occurs at tRNA loci

Non-coding RNAs (ncRNAs) are often decorated with post-transcriptional modifications that render them difficult to amplify using reverse transcriptase ^27,28^. In particular, tRNA derivatives formed by cleavage and base-specific modifications are interesting because they serve functions beyond their canonical role in translation ^29,30^, with implications for pathogenesis ^31,32^ and bacterial physiology ^33,34^. Quantifying microbiome tRNA abundances often requires mass spectrometry or tailored protocols to remove these modifications prior to sequencing ^35,36^. We compared active transcription of tRNA loci in PRO-seq libraries to mature transcripts in RNAseq libraries. We initially focused on three *Prevotella* species found in high abundance in one of the samples for which we could annotate numerous tRNA isoforms. We noticed that a greater proportion of PRO-seq reads could be attributed to these loci than RNAseq reads (0.21 ± 0.07% vs. 0.013 ± 0.016%) and that a larger number of isoforms per tRNA were observed (Supplemental Figure 2), highlighting the utility of PRO-seq to capture differences in ncRNA transcription between closely related bacterial strains.

Among metagenomic tRNA loci, we noticed pile-ups of PRO-seq read 5’-ends within tRNA gene bodies (Figures 4A & B, Supplemental Figure 3), a phenomenon also observed within the cultured *E. coli* heatshock samples (Supplemental Figure 4). We hypothesized that this may be due to processing of tRNAs into tRNA fragments, which act as signaling molecules in many bacterial species ^32^. In one example of a tRNA gene cluster in *Ruminococcus bromii*, we noticed PRO-seq read 5’ end pile-ups in Arg, His and Lys tRNA genes, corresponding to predicted tRNA cleavage sites within each anticodon loop (Figure 4C). Transcription of this locus was absent in the RNAseq data, despite comparable transcription detected across both RNAseq and PRO-seq libraries at protein-coding genes on the same contig (Figure 4D). This example, among others present in a diverse set of species (Supplemental Figure 3), suggests that, similarly to CRISPR loci, tRNA processing is temporally coupled with transcription.

**Figure 4.**
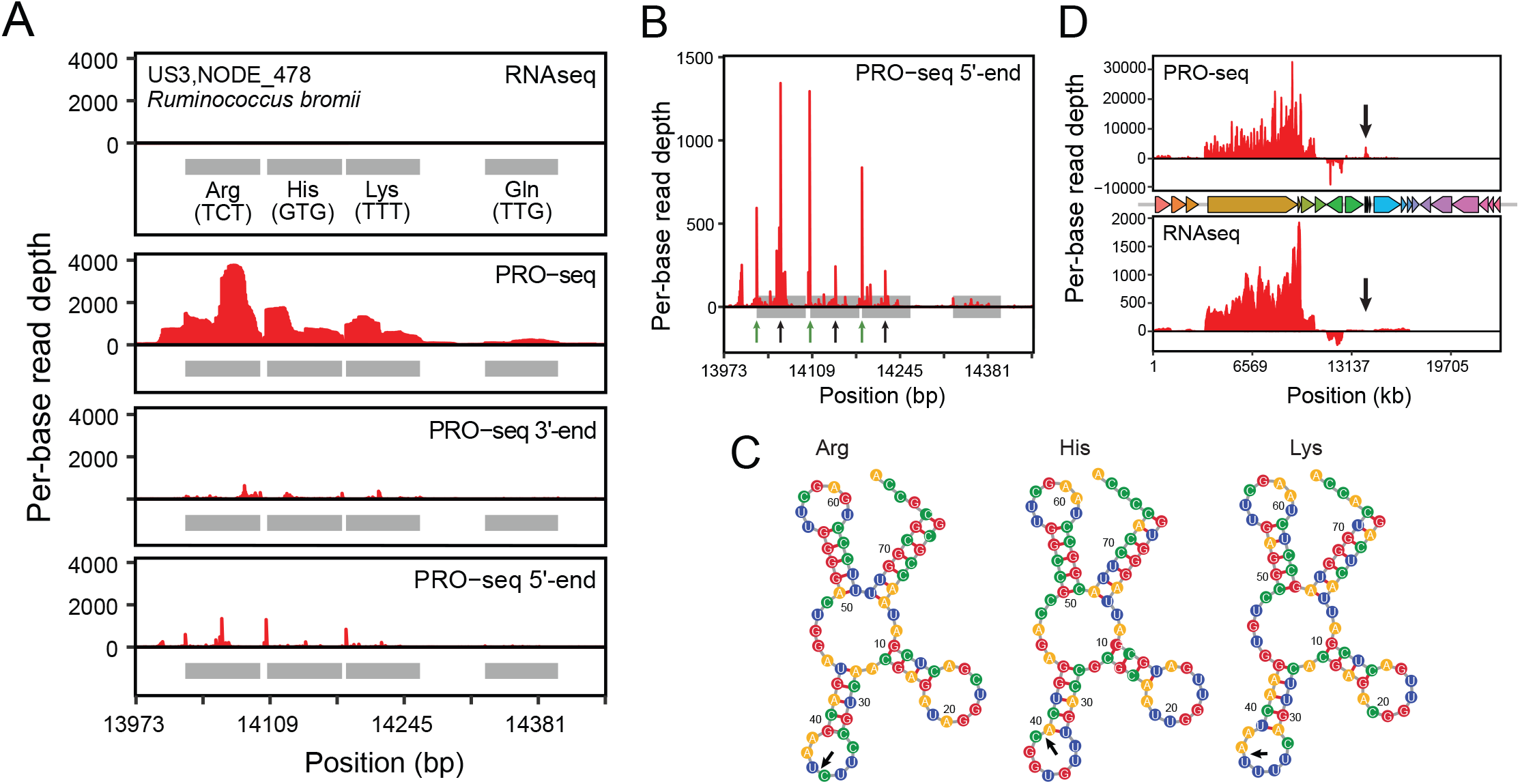
Concurrent transcription and cleavage of tRNAs is observed in PRO-seq libraries. (A) Coverage of PRO-seq and RNA-seq reads across a tRNA gene cluster in a US3 *Ruminococcus bromii* contig. Shaded boxes represent tRNA genes.. Small black arrows in the “PRO-seq 5’ end” panel correspond to the base of the predicted repeat stem loop that serves as the site of crRNA cleavage. (B) Coverage of PRO-seq 5’ ends for the tRNA array shown in (A). Green arrows show the starts of the tRNA genes. Black arrows show the predicted cleavage sites within anticodon loops. (C) Coverage of PRO-seq and RNAseq reads over the entire contig. The positions of gene bodies are shown (middle). Black arrows point to the site of the tRNA array shown in (A). (D) Predicted structures and cleavage sites (black arrows) of tRNA genes shown in (A).

There are at least two alternative hypotheses concerning our interpretation of the pile-up of PRO-seq read 5’ ends within tRNA anticodon loops. (1) In the PRO-seq protocol, nascent RNAs are fragmented by alkaline hydrolysis prior to 3’ adapter ligation. ssRNA is more susceptible to chemical hydrolysis than dsRNA ^37^, so unprotected bases within tRNA loops may be overrepresented as sites for hydrolysis products for a given tRNA isoform. If this is the case, we would expect to see similar peaks in the 5’ end pile-up at the T- and D-arms of nascent tRNAs. However, secondary peaks within PRO-seq traces are small and uncommon relative to peaks coincident with anticodon loops at tRNA midpoints (Supplemental Figure 3), suggesting that preferential hydrolysis of non-base-paired RNA cannot fully explain the patterns we observe. (2) A crucial step in all RNA sequencing protocols is reverse transcription, by which DNA is created from an RNA template for library construction and sequencing. Reverse transcriptase (RT) is sensitive to both the structural conformation and chemical modifications of the template RNA strand ^38,39^, and anticodon loops are common sites for methylation in bacteria ^40,41^. Therefore, tRNA anticodon loops may be a common site for RT stalling, leading to false inference of stall sites as the 5’ ends of nascent transcripts. However, 5’ adapter ligation is carried out *before* reverse transcription as part of the PRO-seq protocol, so cDNAs made from aborted RT products will lack a 5’ adapter and the concomitant PCR handle. It is therefore unlikely that such truncated cDNAs would be represented in the sequencing library, and RT stalling is therefore insufficient to explain the patterns we observe.

### RNAP pause-site motifs annotated in diverse species

The procession of RNAP across a gene body can be interrupted by pauses at specific sequences or secondary structures. These pauses are involved in synchronizing transcription and translation, coordinating the recruitment of regulatory factors, and the dissociation of elongation complexes ^42,43^. Transcription pause sites have previously been shown to differ between *E. coli* and *B. subtilis* ^2,44^, suggesting that they may vary across the members of the gut microbiome. To test this, we called PRO-seq 3’ end peaks across gene bodies in well-covered and near-complete metagenome-assembled genomes. We found concordant consensus pause sites between members of the Bacteroidetes phylum, a *Parabacteroides* and a *Prevotella* species, across two individuals (Figure 5). This TA-rich consensus site is similar to that found in *Agathobacter rectale*, a Firmicutes species found in both individuals’ microbiomes. On the contrary, the pause site motif identified in *Sutterella wadsworthensis*, a Proteobacteria, was more closely aligned with the consensus pause site identified in our earlier experiments with *E. coli*, a different Proteobacteria. These observations suggest that PRO-seq is applicable to study questions of comparative transcription regulation across diverse bacterial species.

**Figure 5.**
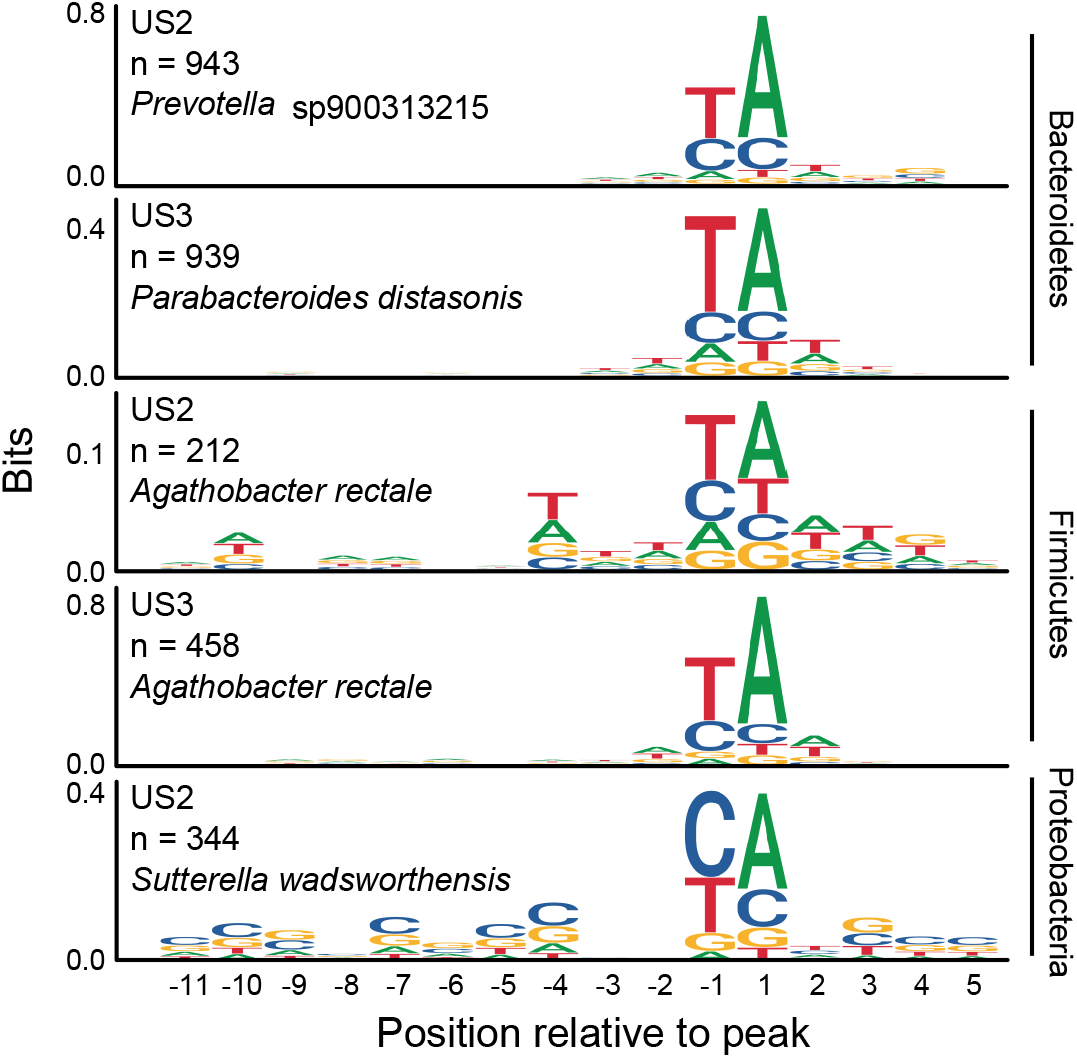
Phyla-specific pause site motifs. Logos for sequences surrounding PRO-seq read 3’ end peaks annotated for two Bacteroidetes, two Firmicutes and one Proteobacteria across two microbiome samples. Position −1 represents the RNAP pause site and position 1 represents the next nucleotide added.

## DISCUSSION

Bacterial nascent transcriptomics provides insights into co-transcriptional RNA processing and RNAP activity and localization. Rather than antibodies or epitopes fused to RNAP, PRO-seq leverages the universal function of RNAP to profile active transcription. We demonstrate its applicability to diverse species within microbiomes, showing that PRO-seq is capable of capturing nascent transcriptional dynamics without the need for cell culture. Our experiments using PRO-seq on heat-shocked *E. coli* highlight the potential for PRO-seq to decipher regulatory circuits operating under other environmental perturbations. Our observation of transcription of RNA species unique to the PRO-seq libraries in both cultured *E. coli* and gut microbiome samples (Figures 3 & 4, Supplemental Figures 1-6) illustrates the utility of PRO-seq in identifying regulatory non-coding RNAs and co-transcriptionally processed RNA products. For bacteria in which non-coding RNAs have not yet been documented, PRO-seq offers a means to broadly survey these RNA molecules. In *E. coli*, where non-coding RNAs are well-annotated, we find that they are enriched in PRO-seq libraries compared to RNAseq libraries (Supplemental Figures 5 & 6). PRO-seq data from metagenomes can therefore provide better guidance on the outputs of these genes than RNAseq data alone. Similarly, given the observation that PRO-seq can be used to broadly profile the transcription of tRNA isoforms and CRISPR loci across diverse species, we expect this method to shed light on the expression and processing of these molecules across conditions.

PRO-seq can also be combined with existing transcriptomic tools to examine transcriptional dynamics at a much finer scale than achievable with RNAseq alone. PRO-seq reveals RNAP positioning with basepair resolution and also captures immature RNA cleavage products. We note that Cappable-seq, the only other nascent transcription method applied to microbiomes, is not suited to the identification of co-transcriptionally processed RNAs due to the need for intact 5’-PPP. However, PRO-seq can also be combined with Cappable-seq for paired analysis of transcription start sites and RNAP localization. PRO-seq may be further paired with NET-seq, albeit in genetically tractable organisms due to its reliance on the immunoprecipitation of RNAP, to discriminate nascent transcripts that are being actively polymerized from those in backtracked states ^45^. Altogether, PRO-seq demonstrates that a larger fraction of bacterial genomes is actively transcribed than represented by traditional RNA sequencing, and that nascent transcription of microbiomes has potential, in concert with other -omics methods, to uncover co-transcriptional dynamics that provide functional insight into the gut microbial community.

## MATERIALS AND METHODS

### *E. coli* heat-shock experiment

An overnight culture of *E. coli* MG1655 was subcultured in 50 mL LB and grown at 37°C to OD600 = 0.95. The culture was then split into 2 × 25 mL, with one half subjected to continued incubation at 37°C and the other half subjected to heat shock at 50 °C for 7 minutes, as described elsewhere ^46^. Cultures were then split into 50 mL aliquots and pelleted by centrifugation at 3000 × g. At this point, pellets were either flash-frozen and stored at −80 °C for RNAseq or carried through permeabilization for PRO-seq.

### Human gut microbiome sample collection

Freshly voided stool samples were collected and homogenized in an equal volume of cold, O2-depleted phosphate-buffered saline, pH 7.2. Stool slurries were centrifuged to remove insoluble material (500 × g, 4 °C, 10 min.), then 12 mL of the liquid supernatant was layered over 3 mL 50% Nycodenz (Accurate Chemical) and centrifuged to concentrate cells (5000 × g, 4 °C, 20 min.). The cell-rich layer above the Nycodenz was collected and stored on ice; this was repeated until all stool homogenate was processed. 2 mL aliquots of each sample were stored at −80 °C for metagenome preparation and RNAseq. Four 600 μL aliquots of each sample were kept on ice until PRO-seq permeabilization. All individuals gave informed consent and all samples were collected under protocol #1609006585 approved by the Cornell University Institutional Review Board.

### RNAseq sample preparation and sequencing

For each *E. coli* treatment, RNA was extracted from replicate samples using the RNeasy Mini Kit (Qiagen), including the optional β-mercaptoethanol treatment specified in the manufacturer’s protocol. On-column DNase I treatment was carried out using components of the DNase Max Kit (Qiagen). RNA was eluted in at least 50 μL nuclease-free water and quantified with the Qubit RNA HS Assay Kit (Thermo Fisher). RNA was combined with 0.1× volume 3M sodium acetate, 3× volumes cold absolute ethanol, and 1 μL GlycoBlue Coprecipitant (Thermo Fisher) and allowed to precipitate at −80 °C for 30 minutes. RNA was pelleted by centrifugation (20,000 × g, 4 °C, 15 min.), washed with cold 70% ethanol, air-dried, and resuspended in nuclease-free water at a concentration of 91 ng/μL. One μg (11 μL) of each sample was subject to rRNA depletion using 2 μL NEBNext Bacterial rRNA Depletion Solution and 2 μL NEBNext Probe Hybridization Buffer (New England Biolabs). Sequencing libraries were prepared from both rRNA-depleted and whole RNA aliquots using the NEBNext Ultra II Directional RNA Library Prep Kit for Illumina (New England Biolabs) following the manufacturer’s protocol for library preparation from intact RNA. RNAclean XP beads were substituted for AMPure XP beads supplemented with 10 U/mL SUPERase-In RNase Inhibitor (Thermo Fisher). Library concentrations were quantified by Qubit dsDNA HS Assay Kit (Thermo Fisher), and library size distributions were visualized by polyacrylamide gel electrophoresis.

For each Nycodenz-purified stool cell sample, RNA was extracted using the RNeasy PowerMicrobiome Kit (Qiagen), following the manufacturer’s protocol to increase the representation of small RNAs. Total RNA was eluted from columns with 100 μL nuclease-free water and quantified using the Qubit RNA BR Assay Kit (Thermo Fisher). Duplicate 1 μg aliquots were subject to RNA fragmentation and rRNA depletion using the QIAseq FastSelect –5S/16S/23S Kit (Qiagen), assuming a RNA integrity number ≥ 8 for all samples. Sequencing libraries were prepared from rRNA-depleted samples as described for *E. coli*. Sequencing platforms and number of reads for each replicate are listed in Supplemental Table 1.

### PRO-seq sample preparation and sequencing

For each *E. coli* treatment, the cell pellets described above were resuspended in 1.5 mL cold cell permeabilization buffer (10 mM Tris-HCl, pH 7.4, 300 mM sucrose, 10 mM KCl, 5 mM MgCl_2_, 1 mM EGTA, 0.05% v/v Tween-20, 0.1% v/v IGEPAL CA-630, 0.1% v/v Triton X-100, 0.5 mM DTT, 1× Roche cOmplete Protease Inhibitor Cocktail (Sigma-Aldrich), and 20 U/mL SUPERase-In RNase Inhibitor (Thermo Fisher); modified from Mahat *et al.* ^8^) and incubated on ice for 5 minutes. Pelleting, resuspension in permeabilization buffer, and incubation was repeated for a total of 3 permeabilization washes. Cell lysates were then pelleted by centrifugation (10,000 × g, 4 °C, 5 min.), resuspended in 250 μL storage buffer (10 mM Tris-HCl, pH 8.0, 25% v/v glycerol, 5 mM MgCl_2_, 0.1 mM EDTA, and 5 mM DTT), split into 5 × 50 μL aliquots, flash-frozen on dry ice / ethanol, and stored at −80 °C until run-on. Final cell concentrations inferred from plating pre-permeabilization cell suspensions were 2.5 × 10^10^ and 5.0 × 10^10^ CFU/mL for control and heat-shocked samples, respectively.

To improve the permeabilization of Gram-positive organisms, 1000 U of Ready-Lyse Lysozyme Solution (Lucigen) was added to each 600 μL Nycodenz-purified stool cell aliquot and incubated for 10 minutes on ice. Then, cell suspensions were transferred to 2 mL screw-cap tubes and combined with 400 μL sterile 0.5 mm glass beads and 1 mL cold cell permeabilization buffer. Cells were pulverized by vortexing for 3 cycles of 2 minutes at max Hz followed by 2 minutes on ice. Lysates were stored upright on ice for 10 minutes to allow beads to settle, then 1 mL supernatant from each tube was transferred to a 1.5 mL tube and centrifuged to collect cell contents (10,000 × g, 4 °C, 5 min.). Pellets were washed once with 1 mL cold storage buffer, pelleted again, and resuspended in 200 μL cold storage buffer. Lysates were flash-frozen and stored as described for *E. coli*.

For all samples, PRO-seq was carried out following the “4-Biotin run-on” variant of the protocol described in Mahat *et al* ^8^. Briefly, permeabilized cells were thawed on ice and run-on reactions were carried out at 37 °C using a master mix containing Biotin-11-ATP, Biotin-11-CTP, Biotin-11-GTP, and Biotin-11-UTP. Total RNA was extracted by TRIzol and ethanol precipitation, and RNA was fragmented by NaOH hydrolysis. 3’ adapters were ligated, then biotinylated transcripts were enriched and washed with hydrophilic streptavidin magnetic beads. 5’ de-capping and phosphorylation were carried out with nascent transcripts bound to the beads, then RNA was eluted from the beads by TRIzol extraction and ethanol precipitation. 5’ adapters with unique molecular identifiers were ligated to nascent transcripts, and excess adapters were removed by again capturing biotinylated RNA on streptavidin beads, washing the beads, and re-extracting RNA with TRIzol and ethanol precipitation. Nascent RNA was reverse transcribed, and cDNA was quantified by qPCR to determine the appropriate number of cycles for PCR amplification. Library amplification was carried out using custom PCR primers to incorporate Illumina adapter sequences and i7 barcodes. PCRs were cleaned up with Exonuclease I and Shrimp Alkaline Phosphatase. DNA concentration was quantified with the Qubit dsDNA HS Assay Kit, and library quality was assessed by polyacrylamide gel electrophoresis. Sequencing platforms and number of reads for each replicate are listed in Supplemental Table 1.

### Metagenomic library preparation, sequencing, assembly, binning, and annotation

To prepare metagenomes against which to map transcriptomics reads, DNA was isolated from 250 μL of Nycodenz-purified stool cells using the DNeasy PowerSoil Kit (Qiagen). DNA was eluted in 100 μL warm 0.1× TE, quantified by Qubit dsDNA BR Assay, and diluted to 0.2 ng/μL for input to the Nextera XT DNA Library Preparation Kit (Illumina). Sequencing libraries were prepared from 1 ng fecal DNA following the manufacturer’s protocol, and libraries were cleaned up using 1.5× volumes of AMPure XP beads. Library concentration was quantified by Qubit dsDNA HS Assay, and fragment size distribution was visualized by 8% polyacrylamide gel electrophoresis.

Metagenomes were sequenced as referenced in Supplemental Table 1. Raw reads were processed with PRINSEQ lite v0.20.4 ^47^ and trimmomatic v0.36 ^48^ to remove duplicates and sequencing adapters. Reads mapping to the human genome were discarded using BMTagger ^49^. Clean reads were assembled using SPAdes v3.14.0 ^50^ (paired-end mode and --meta option) and reads were aligned to contigs using BWA-MEM v0.7.17 ^51,52^. Contigs were binned using CONCOCT v1.1.0 ^53^, metaBAT v2.12.1 ^54^, and MaxBin v2.2.4 ^55^, then bins from different programs were resolved into metagenome-assembled genomes (MAGs) using DAS Tool v1.1.2 ^56^ with DIAMOND v2.0.4 ^57^ for single copy gene identification. The completeness and contamination of MAGs was assessed with CheckM v1.1.2 ^58^ and taxonomic classifications were assigned to MAGs using GTDB-Tk v1.0.2 ^59^. MAG features were annotated using prokka v1.14.5 ^60^ (-metagenome, --rfam), which uses Prodigal ^61^, ARAGORN ^62^, barrnap ^63^, and Infernal ^64^ for identification of protein-coding sequences, tRNAs, rRNAs, and ncRNAs, respectively.

### Transcriptomics data processing and analysis

PRO-seq reads were processed with proseq2.0.bsh (https://github.com/Danko-Lab/proseq2.0) to trim by quality, remove adapter sequences, and remove duplicates by their unique molecular identifiers (UMIs). RNAseq reads were similarly processed, but without UMI deduplication. Cleaned paired-end reads were aligned to their respective references using BWA: metatranscriptome reads were aligned to the assemblies described above; *E. coli* reads were aligned to the GenBank Reference Sequence for E. coli K12, version NC_000913.3 ^65,66^. Bam files were filtered with SAMtools v1.11 ^67^ to include only paired reads in proper pairs with a minimum MAPQ score of 30 (-f 3 -q 30) and exclude all unmapped or nonprimary alignments (-F 2316). Reads were assigned to features using the featureCounts function from subread v2.0.2 ^68^. *E. coli* protein-coding genes and regulatory loci were identified using the regutools R package ^69^ and RegulonDB v10.9 ^70^. The genomecov function from BEDTools v2.29.2 ^71^ was used to report strand-specific PRO-seq and RNAseq depth at each position in the metagenome (-ibam -d -pc - strand). For PRO-seq, metagenomic depth profiles from 3’ and 5’ fragment ends were additionally reported as follows: since the P5 Illumina adapter is ligated to the 3’ end of the nascent transcript, the 5’ end of the first read in each pair gives the 3’ end of the nascent transcript on the opposite strand (samtools view -f 64 -b $bam | bedtools genomecov −5 -d -strand - > plus_3p.txt); likewise, the 5’ end of the second read in each pair gives the 5’ end of the nascent transcript on the same strand, since proper pairs align to opposite strands (samtools view -f 128 -b $bam | bedtools genomecov −5 -d -strand + > plus_5p.txt).

Read depth profiles at regions of interest were visualized with ggplot2 ^72^ using custom R code available at https://github.com/britolab/PRO-seq. CRISPR repeats were detected using MinCED ^73^, which is derived from CRISPR Recognition Tool ^74^. CRISPR RNA and tRNA secondary structures were predicted with the ViennaRNA secondary structure server ^75^ and visualized with forna ^76^. Pearson’s correlation coefficients (*r*) and Spearman’s rank correlation coefficients (*ρ*) were calculated for correlation plots using the stats package from base R ^77^. Wilcoxon signed-rank tests were performed using the ggpubr package v0.4.0 ^78^. Peaks were called from 3’ end depth data by first filtering all positions for a minimum depth of 10 reads. Then, the mean coverage over a ±25 nt interval surrounding each position was calculated, and Z scores were determined for each peak centered in its interval. Positions with Z scores of at least 5 were kept, and sequences surrounding those peaks were pulled to create sequence logos with the ggseqlogo package ^79^.

## Code and data availability

Scripts and R workbooks are available at https://github.com/britolab/PRO-seq. Sequencing data has been uploaded to NCBI’s Sequence Read Archive and is associated with BioProjects PRJNA800038 and PRJNA800070.

## SUPPLEMENTARY DATA

**Supplemental Figure 1.**
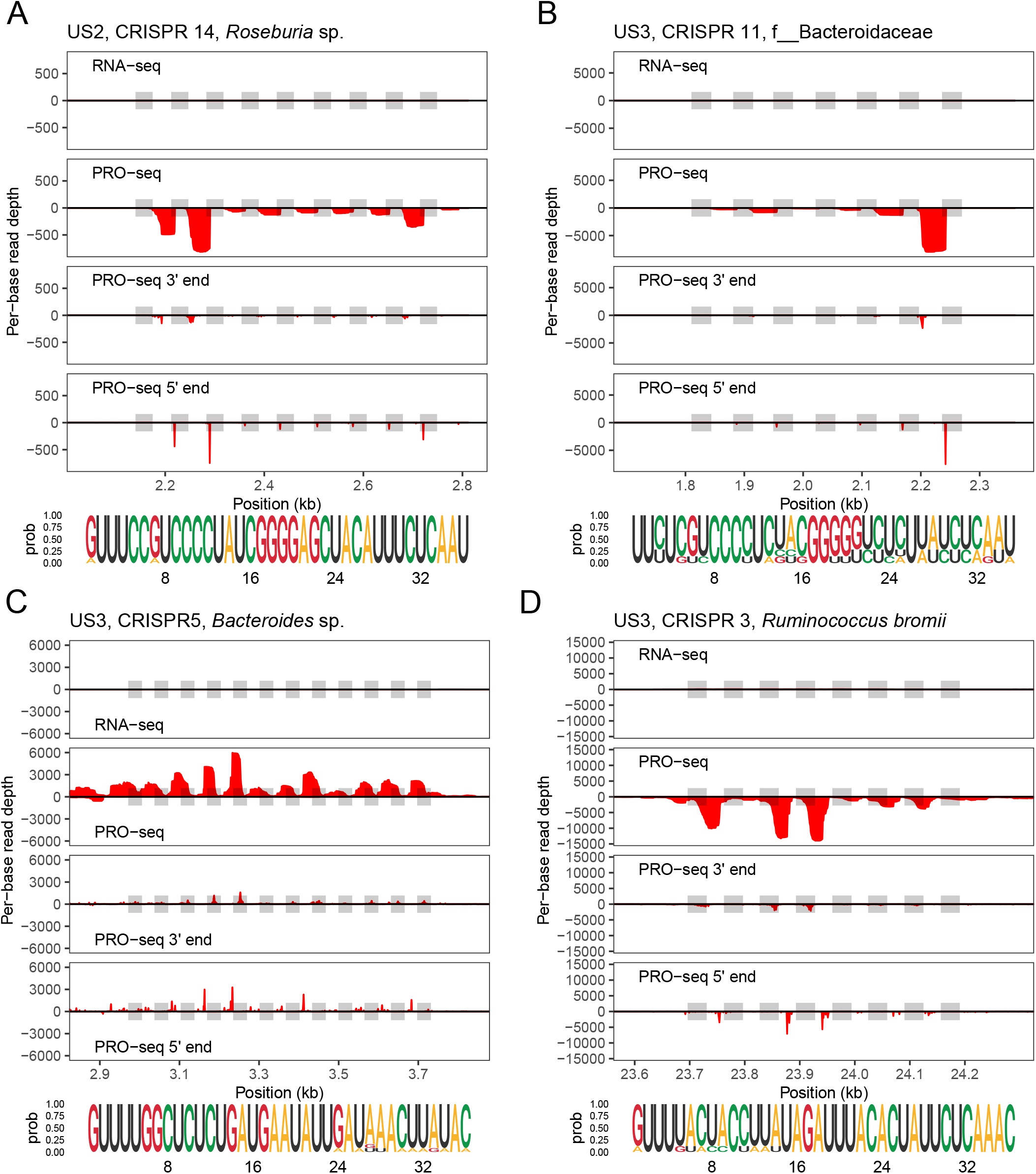
Periodicity observed in CRISPR loci within the PRO-seq data. Strand-specific RNAseq and PRO-seq read depths, in addition to PRO-seq reads’ 3’- and 5’-ends, are plotted for several well-covered CRISPR loci. Shaded boxes represent repeats. Sequence logos below each plot show repeat conservation. As in Figure 3, (A) and (B) show PRO-seq read 5’ end pile-ups at the same position across repeats. (C) and (D) show PRO-seq read 5’ end pile-ups within spacers.

**Supplemental Figure 2.**
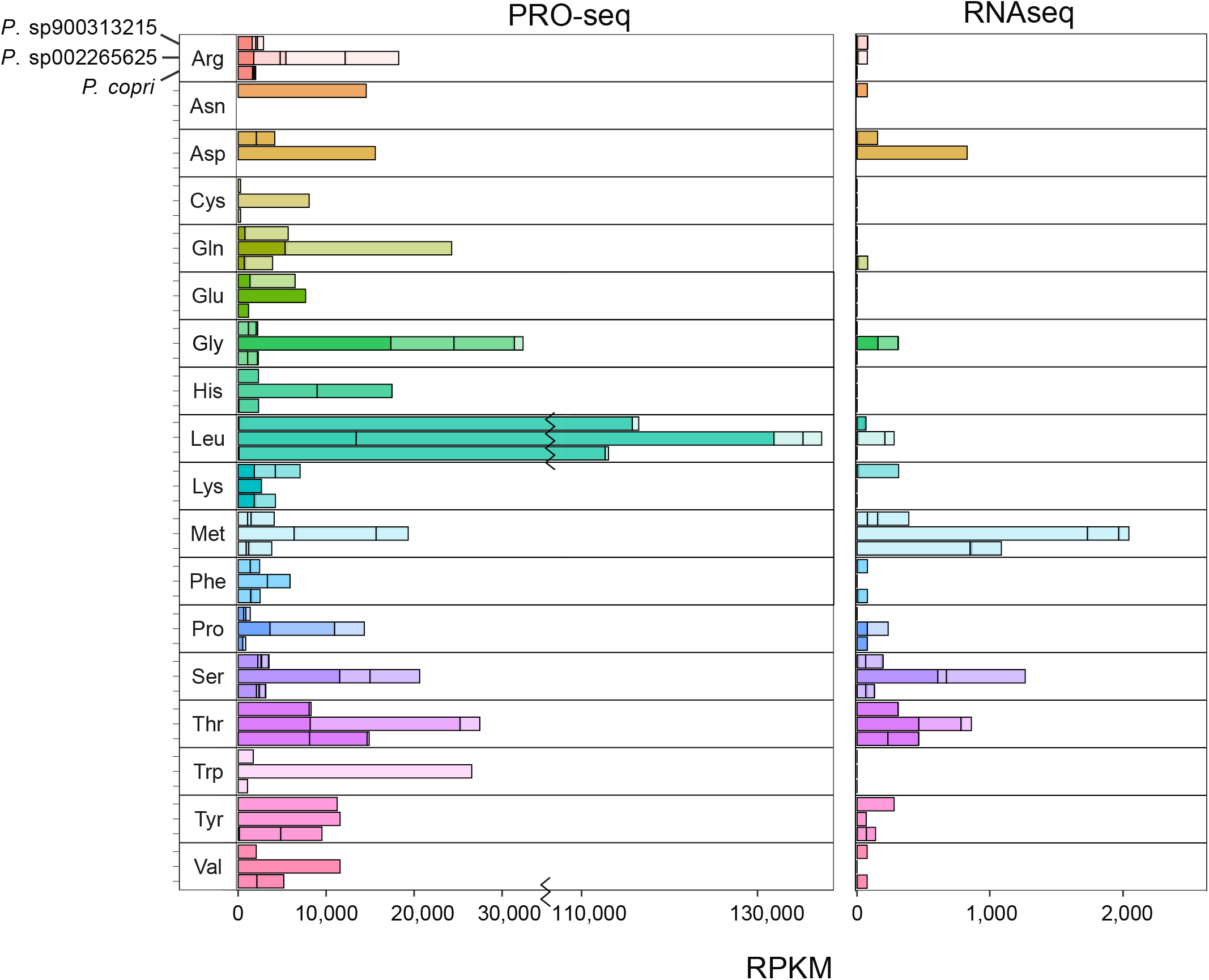
Isoforms of tRNAs in *Prevotella* species are better represented in PRO-seq than RNAseq data. tRNA genes were identified in three highly complete US2 bins: *Prevotella* sp900313215, *Prevotella* sp002265625 and *Prevotella copri*. Different colors in the stacked bar plots represent different tRNA isoforms.

**Supplemental Figure 3.**
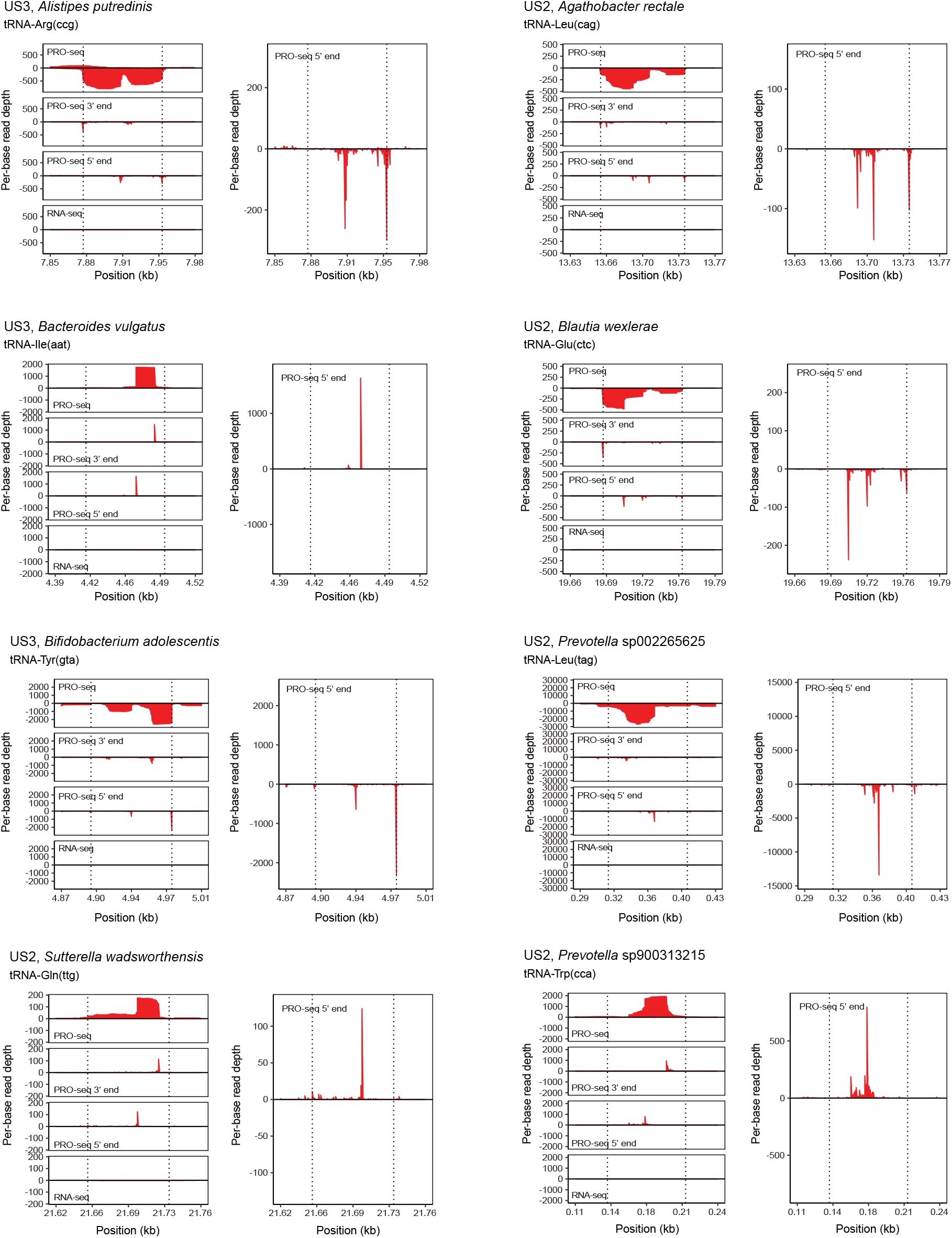
PRO-seq traces showing 5’ read end pile-ups within microbiota tRNA genes. Representative tRNA genes, listed according to the sample, species annotation, and anticodon, are depicted from the two human microbiome samples. PRO-seq coverage, pile-up of PRO-seq 3’and 5’ read ends, and RNAseq coverage are shown for each tRNA gene (left). A zoomed-in PRO-seq read 5’ end pile-up is shown for each tRNA gene (right).

**Supplemental Figure 4.**
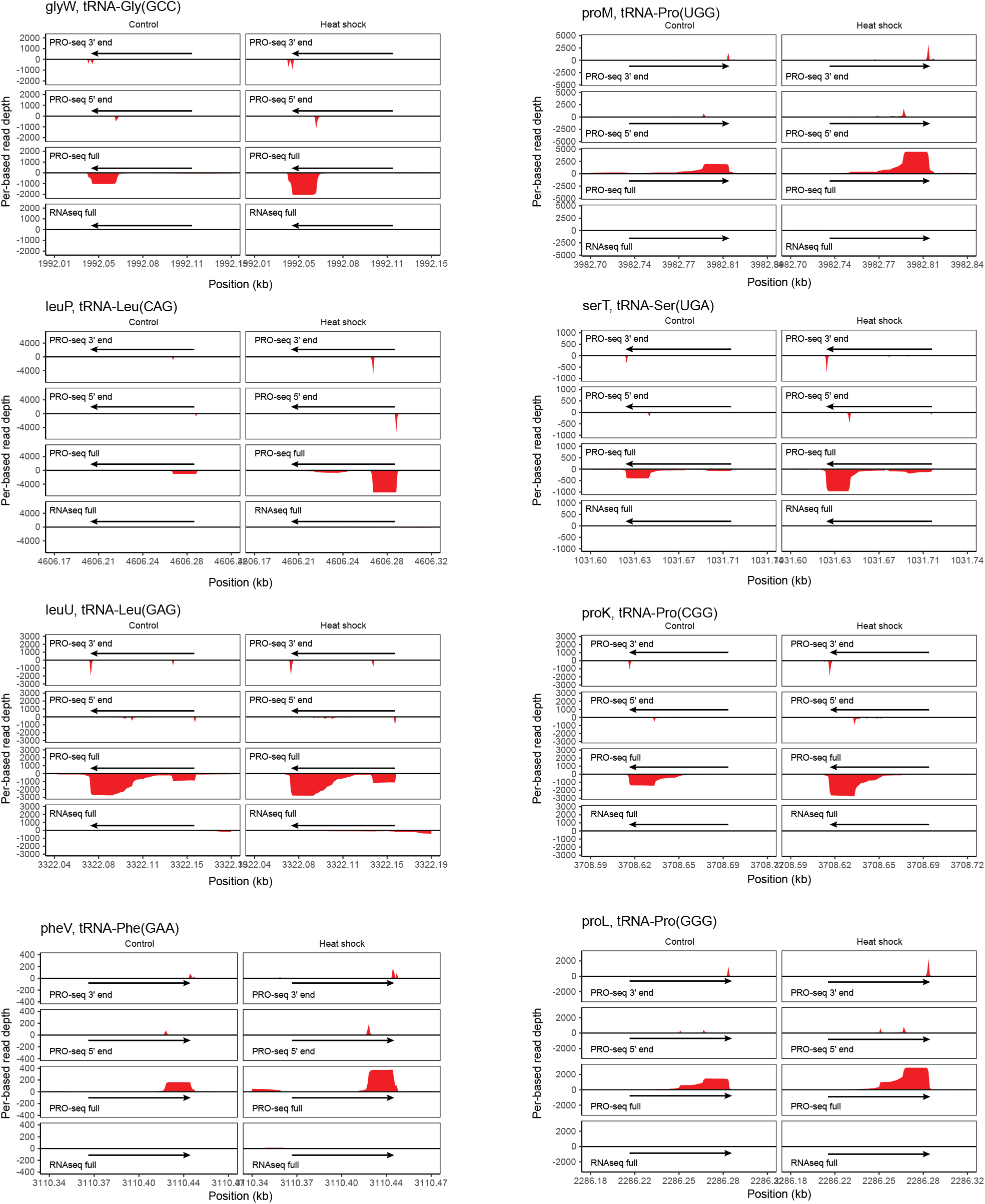
PRO-seq traces across *E. coli* tRNAs show PRO-seq 5’ read end pile-ups. Representative *E.* coli tRNA genes, listed by isoform, are shown for control (left) and heat shock (right) conditions. PRO-seq coverage, pile-up of PRO-seq 3’and 5’ read ends, and RNAseq coverage are shown for each tRNA gene.

**Supplemental Figure 5.**
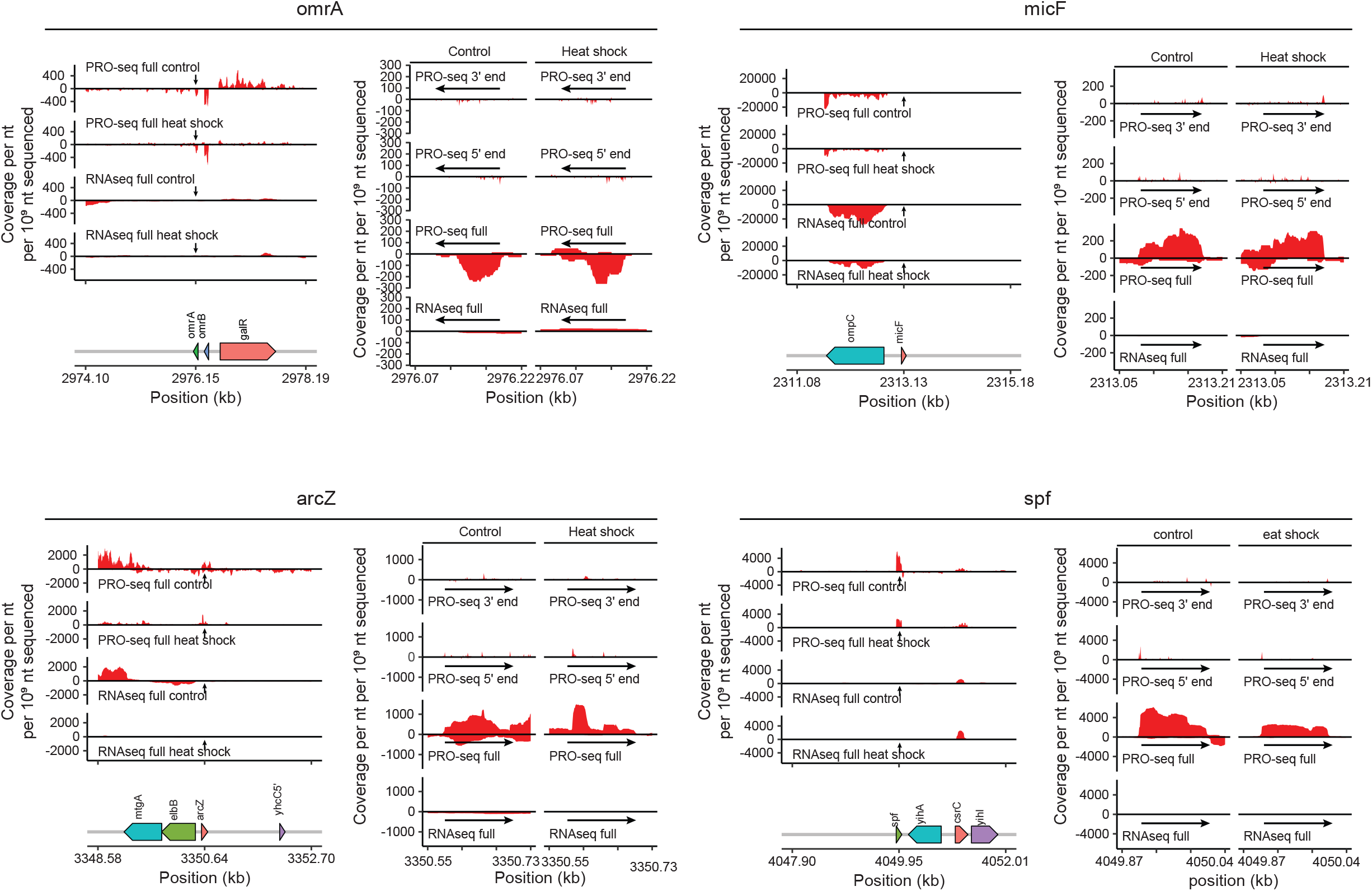
PRO-seq traces capture transcription of *E. coli* small regulatory RNAs. Selected *E. coli* small non-coding RNA (sRNA) loci shown with coverage (per nucleotide per 10^9^ sequenced reads) surrounding each locus (left) in RNAseq libraries and PRO-seq libraries from under control and heat shock conditions. On the right, RNAseq coverage, composite PRO-seq read coverage, 5’ end and 3’ end coverage are shown for the specific portion of the locus encoding the sRNA.

**Supplemental Figure 6.**
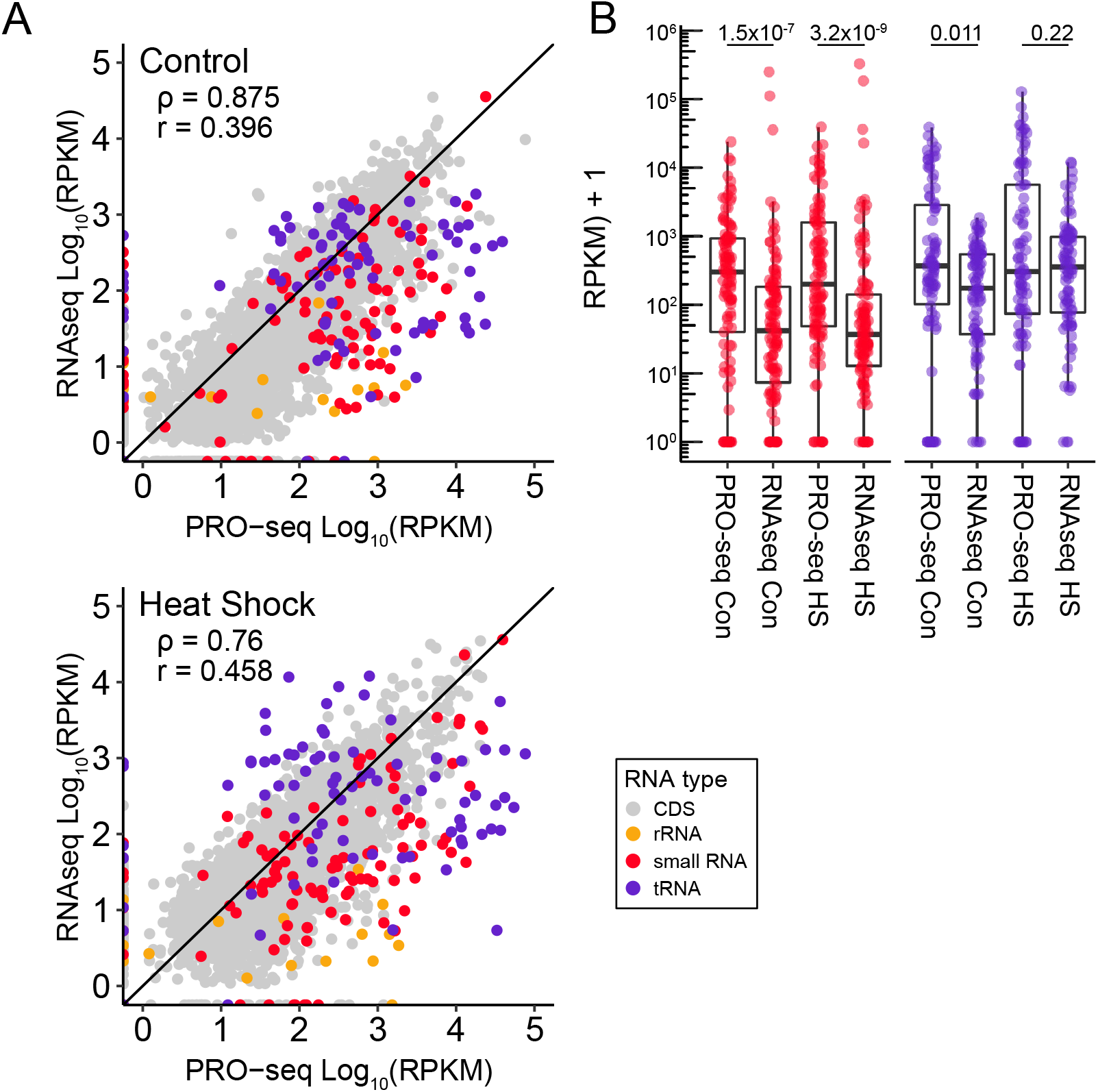
*E. coli* genome-wide detection of noncoding RNAs in PRO-seq versus RNAseq libraries. (A) Log-log RPKM plots comparing merged PRO-seq and RNAseq libraries for control and heatshock conditions. Genes are colored by RNA type. Spearman’s rank correlation coefficients (*ρ*) and Pearson’s correlation coefficients (*r*) are inset. (B) Box plots show the RPKM distribution for small non-coding RNAs and tRNAs across control and heat-shock conditions. Black lines represent medians. P-values from Wilcoxon signed-rank tests are reported for each RNA type + treatment pair.

**Supplemental Table 1.** List of samples sequenced in this project.

| Sample name | Replicate | Sequencing platform | Number of clean paired-end reads |
| --- | --- | --- | --- |
| *E. coli* – control – RNAseq | 1 | NextSeq, 2 × 150 | 1,713,282 |
|  | 2 | NextSeq, 2 × 150 | 1,871,542 |
| *E. coli* – control – RNAseq with rRNA depletion | 1 | NextSeq, 2 × 150 | 1,834,682 |
|  | 2 | NextSeq, 2 × 150 | 1,456,787 |
| *E. coli* – control – PRO-seq | 1 | NextSeq, 2 × 75 | 3,982,364 |
|  | 2 | NextSeq, 2 × 75 | 1,615,517 |
|  | 3 | NextSeq, 2 × 75 | 6,026,982 |
| *E. coli* – heat shock – RNAseq | 1 | NextSeq, 2 × 150 | 1,533,500 |
|  | 2 | NextSeq, 2 × 150 | 1,790,382 |
| *E. coli* – heat shock – RNAseq with rRNA depletion | 1 | NextSeq, 2 × 150 | 1,290,188 |
|  | 2 | NextSeq, 2 × 150 | 1,442,903 |
| *E. coli* – heat shock – PRO-seq | 1 | NextSeq, 2 × 75 | 4,857,572 |
|  | 2 | NextSeq, 2 × 75 | 5,730,267 |
|  | 3 | NextSeq, 2 × 75 | 2,703,945 |
| US2 – RNAseq with rRNA depletion | 1 | NextSeq, 2 × 150 | 23,115,457 |
|  | 2 | NextSeq, 2 × 150 | 19,572,778 |
| US2 – PRO-seq | 1 | HiSeq X, 2 × 150 | 120,441,062 |
|  | 2 | HiSeq X, 2 × 150 | 89,204,805 |
| US3 – RNAseq with rRNA depletion | 1 | NextSeq, 2 × 150 | 21,299,806 |
|  | 2 | NextSeq, 2 × 150 | 21,617,152 |
| US3 – PRO-seq | 1 | HiSeq X, 2 × 150 | 125,455,387 |
|  | 2 | HiSeq X, 2 × 150 | 117,270,406 |
| US2 metagenome | n/a | NextSeq, 2 × 150 | 15,572,009 |
| US3 metagenome | n/a | NextSeq, 2 × 150 | 16,747,211 |

**Supplemental Table 1. List of samples sequenced in this project.**
| **Sample name** | **Replicate** | **Sequencing platform** | **Number of clean paired-end reads** |
| --- | --- | --- | --- |
| *E. coli* – control – RNAseq | 1 | NextSeq, 2 × 150 | 1,713,282 |
|  | 2 | NextSeq, 2 × 150 | 1,871,542 |
| *E. coli* – control – RNAseq with rRNA depletion | 1 | NextSeq, 2 × 150 | 1,834,682 |
|  | 2 | NextSeq, 2 × 150 | 1,456,787 |
| *E. coli* – control – PRO-seq | 1 | NextSeq, 2 × 75 | 3,982,364 |
|  | 2 | NextSeq, 2 × 75 | 1,615,517 |
|  | 3 | NextSeq, 2 × 75 | 6,026,982 |
| *E. coli* – heat shock – RNAseq | 1 | NextSeq, 2 × 150 | 1,533,500 |
|  | 2 | NextSeq, 2 × 150 | 1,790,382 |
| *E. coli* – heat shock – RNAseq with rRNA depletion | 1 | NextSeq, 2 × 150 | 1,290,188 |
|  | 2 | NextSeq, 2 × 150 | 1,442,903 |
| *E. coli* – heat shock – PRO-seq | 1 | NextSeq, 2 × 75 | 4,857,572 |
|  | 2 | NextSeq, 2 × 75 | 5,730,267 |
|  | 3 | NextSeq, 2 × 75 | 2,703,945 |
| US2 – RNAseq with rRNA depletion | 1 | NextSeq, 2 × 150 | 23,115,457 |
|  | 2 | NextSeq, 2 × 150 | 19,572,778 |
| US2 – PRO-seq | 1 | HiSeq X, 2 × 150 | 120,441,062 |
|  | 2 | HiSeq X, 2 × 150 | 89,204,805 |
| US3 – RNAseq with rRNA depletion | 1 | NextSeq, 2 × 150 | 21,299,806 |
|  | 2 | NextSeq, 2 × 150 | 21,617,152 |
| US3 – PRO-seq | 1 | HiSeq X, 2 × 150 | 125,455,387 |
|  | 2 | HiSeq X, 2 × 150 | 117,270,406 |
| US2 metagenome | n/a | NextSeq, 2 × 150 | 15,572,009 |
| US3 metagenome | n/a | NextSeq, 2 × 150 | 16,747,211 |

## FUNDING

Funding was provided by the Genomics Innovation Hub and the Center for Vertebrate Genomics (CVG) at Cornell University, the National Institutes of Health (1DP2HL141007). A.V. is a CVG Distinguished Scholar. I.L.B. is a Sloan Foundation Research Fellow, a Pew Scholar in the Biomedical Sciences and a Packard Fellow for Science and Engineering.

## CONFLICT OF INTEREST

The authors have no conflicts of interest to declare.

## Notes

### Competing Interest Statement

The authors have declared no competing interest.

